# *Trypanosoma brucei gambiense* group 2 experimental *in vivo* life cycle: from procyclic to bloodstream form

**DOI:** 10.1101/2023.10.19.562698

**Authors:** Paola Juban, Jean-Mathieu Bart, Adeline Segard, Vincent Jamonneau, Sophie Ravel

## Abstract

*Trypanosoma brucei gambiense* (*Tbg*) group 2 is a subgroup of trypanosomes able to infect humans in West and Central Africa. Unlike other agents causing sleeping sickness such as *Tbg* group 1 and *Trypanosoma brucei rhodesiense, Tbg*2 lacks the typical molecular markers associated with resistance to human serum. Only thirty-six strains of *Tbg*2 have been documented, and therefore, very limited research has been conducted despite its zoonotic nature. Some of these strains are only available in their procyclic form which hinders human serum resistance assays and mechanistic studies. Furthermore, the understanding of *Tbg*2’s potential to infect tsetse flies and mammalian hosts is limited. In this study, 165 *Glossina palpalis gambiensis* flies were experimentally infected with procyclic *Tbg*2 parasites. 35 days post-infection, 43 flies out of the 80 still alive flies were found *Tbg2* PCR-positive in the saliva. These flies were able to infect 3 out of the 4 mice used for blood-feeding. Dissection revealed that only six flies really carried mature infections in their midguts and salivary glands. Importantly, a single fly with a mature infection was sufficient to infect a mammalian host. This *Tbg2* transmission success confirms that *Tbg*2 strains can establish in tsetse flies and infect mammalian hosts. The study describes an effective *in vivo* protocol for transforming *Tbg*2 from procyclic to bloodstream form, reproducing the complete *Tbg*2 cycle from *G. p. gambiensis* to mice. These findings provide valuable insights into *Tbg*2’s host infectivity, and will facilitate further research on mechanisms of human serum resistance.

## Introduction

*Trypanosoma brucei* (*Tb*) is an extracellular protozoan parasite transmitted by an arthropod hematophagous vector: the tsetse fly (*Glossina sp*) (Hoare, 1972). Among the *Tb* species, the *Tb brucei* (*Tbb*) sub-species causes animal African trypanosomiasis or nagana in fauna. The human form of the disease, human African trypanosomiasis (HAT) or sleeping sickness is caused by the two other *Tb* sub-species: *Tb rhodesiense* (*Tbr*) and *Tb gambiense* (*Tbg*) (Büscher et al., 2017). *Tbr* causes an acute form of the disease in East Africa whereas *Tbg* develops into a chronic form in Central and West Africa. *Tbg* HAT was responsible of 87% of the reported cases in 2019-2020 and is targeted by World Health Organization for the interruption of transmission by 2030 (Franco et al., 2022).

During the 1990’s, the development of new analytical molecular methods allowed the division of the *Tbg* subspecies in two groups. The most prevalent, genetically homogenous and monophyletic was the group 1 (*Tbg*1) (Gibson, 1986), invariably resistant to normal human serum (NHS) particularly thanks to the expression of the *Tbg-*specific glycoprotein (TgsGP) (Berberof et al., 2001; Uzureau et al., 2013). In a recent review, group 2 (*Tbg*2) was defined as all human-infective *Tb* trypanosomes from West and Central Africa that do not fit into *Tbg*1 using various molecular markers (Jamonneau et al., 2019). *Tbg*2, *Tbb* and *Tbr* are obviously highly diverse lineages but *Tbg*2 is different from *Tbr* with a consistent lack of serum resistance associated gene (SRA) (Gibson et al., 2002). If *Tbg*2 does not appear to be a public health problem with only thirty-six strains referenced in the literature regarding the above definition, it represents a zoonotic form of HAT with a risk of transmission from animals to humans. In the current elimination context, it seems crucial to be able to detect such infections using adapted effective diagnosis and to determine if they are due to human serum resistance (HSR) trypanosomes or patient immunodeficiency (constitutive or transient) in order to implement adapted control strategies. *Tbg2* stocks from different labs were gathered at UMR INTERTRYP (IRD/CIRAD, Montpellier, France) to study the HSR mechanisms and provide essential elements to anticipate the appearance of new mechanisms and to prevent possible phenomena of emergence (Jamonneau et al., 2019).

*Tb* parasites have a multistage life cycle divided between the tsetse fly vector and a mammalian host. Along this life cycle, the parasite should continuously adapt to its surrounding environment. In the mammalian host, the bloodstream form (BSF) trypanosomes either exists in a proliferative long slender form or in a quiescent short stumpy form pre-adapted to the vector (Figure 1). Following the infective blood meal, trypanosomes transform into their replicative procyclic form (PCF) in the tsetse fly’s midgut. Approximately one month after the infective meal, a small proportion of tsetse flies (about 0.01% in natural conditions) colonizes the salivary glands where they attach as epimastigote forms (EMF) (Frézil et al., 1994). Trypanosomes finally differentiate into infectious metacyclic forms (MCF) which can be transmitted to the mammalian host during the next blood meal.

**Figure 1:**
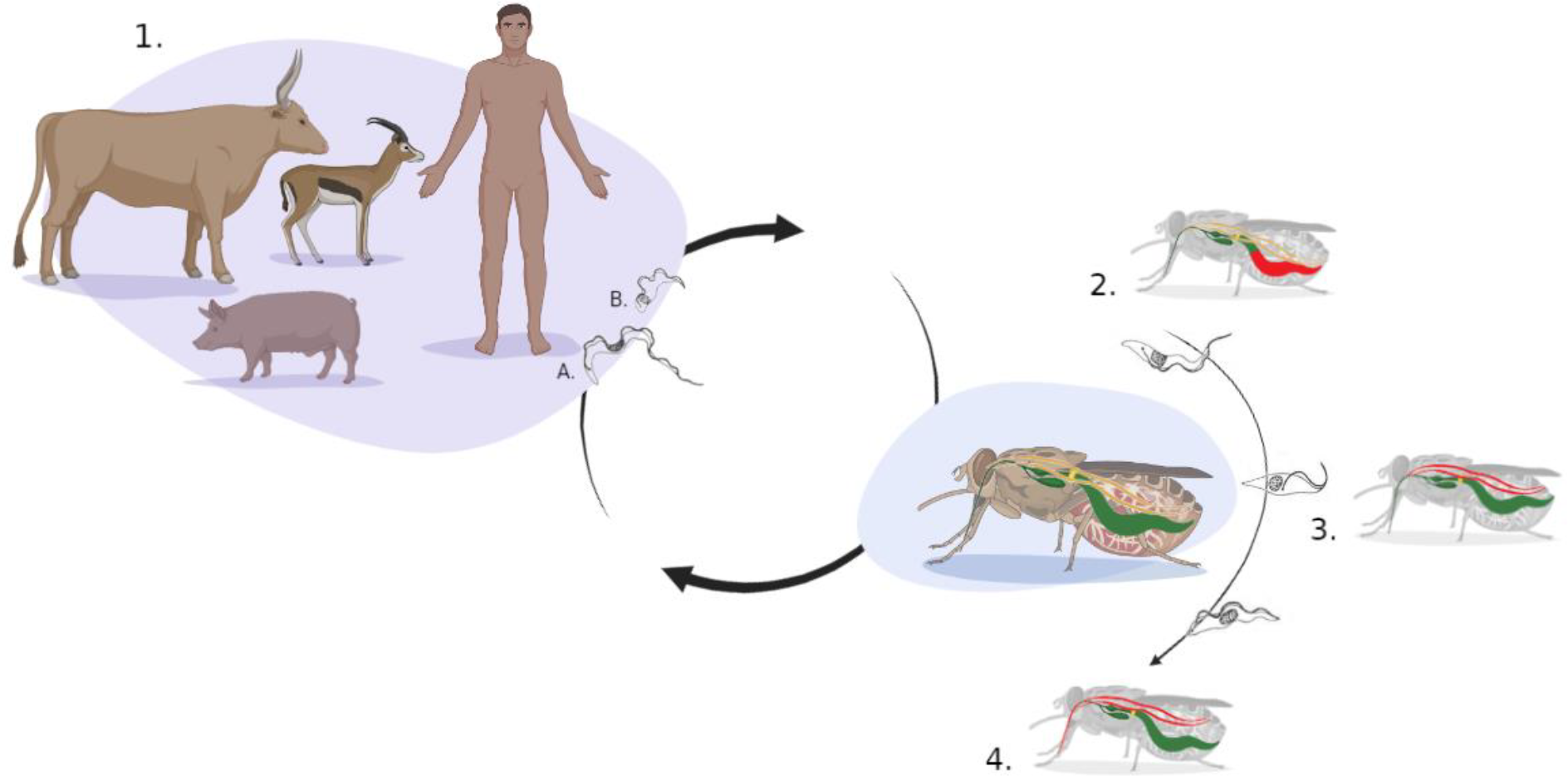
Schematic life cycle of *Trypanosoma brucei*. 1: Bloodstream trypanosomes in mammalian host A: long slender form. B: short stumpy form. 2: Procyclic trypanosomes in tsetse fly’s midgut. 3: Epimastigote trypanosomes colonize salivary glands. 4: Metacyclic trypanosomes in salivary glands can be transmitted to a mammalian host during the next tsetse fly meal. In red: localization of the parasites into the tsetse fly through the cycle.

Most of the *Tbg*2 strains are only available in their PCF and cannot be tested for their resistance to NHS and for the study of the mechanisms implied. Moreover, very few is known about the potential for infection of *Tbg*2 in tsetse and mammalian host. Some rare studies have been conducted using PCF of *Tbg*1 or *Tbg*2 but transmissions to a mammalian host were not successful or were not attempted (Richner et al., 1988, Ravel et al., 2003). Very few experimental studies of the transmission cycle of *Tbg* have been conducted because of its low rate of mature infections (Maudlin and Welburn, 1994). To our knowledge, a complete transmission cycle of *Tbg* under experimental conditions has never been documented. The objectives of the present experimental study were (i) to confirm that *Tbg*2 PCF can settle in tsetse flies and become infectious to mice and (ii) to transform *Tbg*2 PCF to BSF for further studies on HSR.

## Material and Methods

### Tsetse flies

In this study, tsetse flies were used from a colony of *Glossina palpalis gambiensis* originated from Burkina Faso and maintained at CIRAD (Montpellier, France). Only males were chosen, as they develop a higher proportion of salivary glands infections (mature infections) compared to females when experimentally infected with trypanosomes of the subgenus *Trypanozoon* (Maudlin et al., 1991, Dale et al., 1995, Milligan et al., 1995). Due to several issues relative to fly physiology (flies’ natural death, fluctuating infecting meal feedings, low rate of trypanosomes colonization in the midgut and of mature infection in the salivary glands) already described elsewhere (Moloo et al., 1986; Maudlin and Welburn, 1994), 165 tsetse flies were used assuming that this number would be sufficient to obtain mature infections after one month. Before the infection process, one wing was removed from each fly for security reason.

### Trypanosomes

The *Tbg*2 HTAG107-1 strain, also known as IPR107-1, was used in this study. This strain was isolated from human in 1986 in the Daloa HAT focus, Côte d’Ivoire (Stevens et al., 1992). HTAG107-1 was grown in SDM-79 medium (Brun et al., 1979) supplemented with 20% fetal calf serum previously decomplemented (30 minutes at 56°C). PCF trypanosomes were cultivated at 28°C and harvested when 2.4×10^8^ parasites were obtained (1×10^7^ p/mL) (Figure 2). Parasites were then collected after centrifugation (1500×g, 10minutes) and the pellet containing trypanosomes was washed three times with phosphate-buffered saline (PBS 1X) before use.

**Figure 2:**
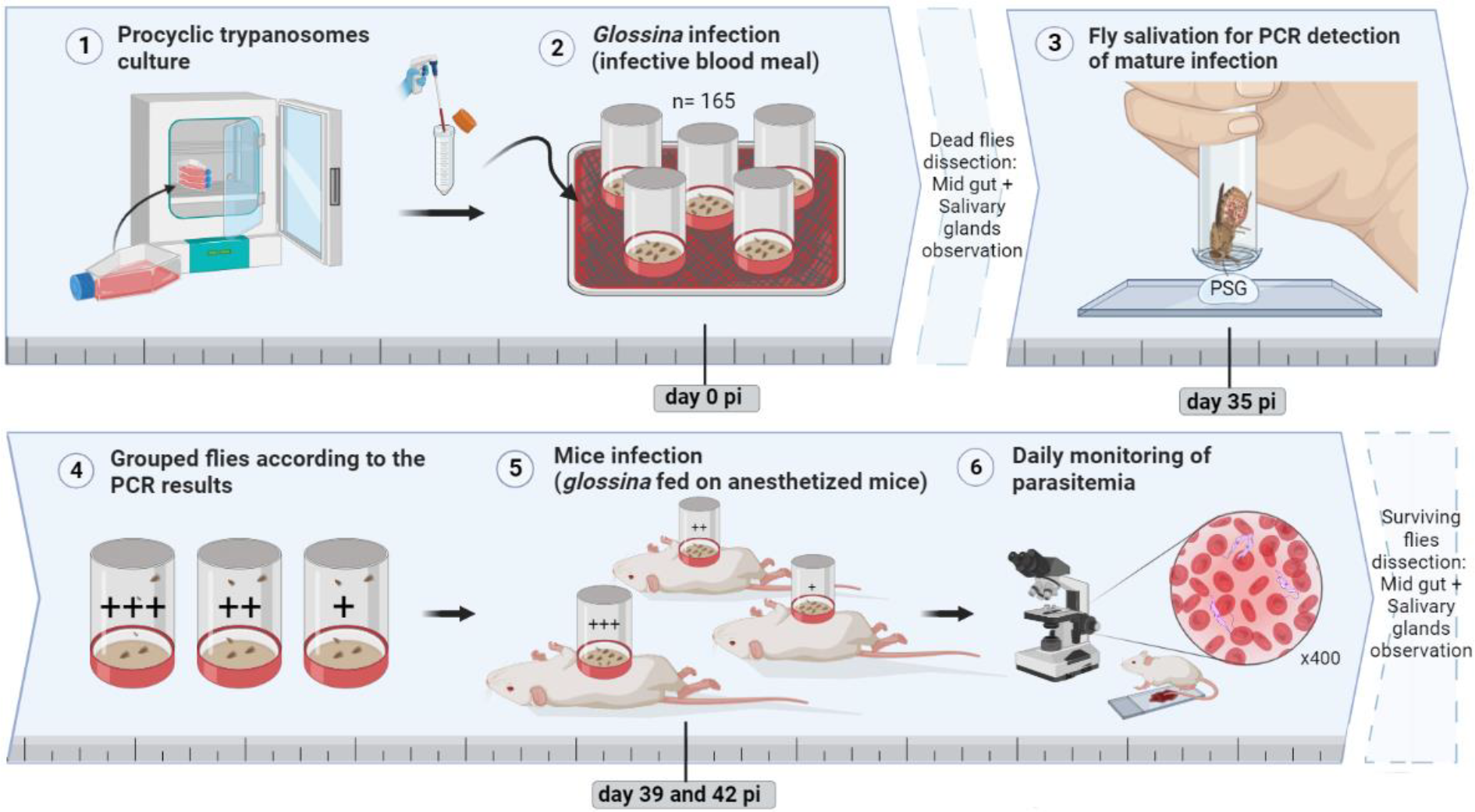
*Tbg*2 experimental in *vivo* cycle protocol (pi: post infection). +++: strong PCR signal; ++: medium PCR signal; +: weak PCR signal

### Experimental infection of tsetse flies

PCF trypanosomes (2.4×10^8^) were gently mixed with 40 ml of sheep blood heated to 31°C. The infected blood was proposed to starved *G. p. gambiensis* teneral males through a silicone membrane (Bauer and Wetsel, 1976) (Figure 2). After infective meal, tsetse flies were separated according to their blood-feeder status (blood meal visible in the abdomen or not). After 24h, the process of infective meal was repeated with the non-gorged tsetse flies to ensure a maximum infection rate success. Flies were then fed with uninfected sheep blood, three days a week for 35 days.

### Salivation assay and PCR

Thirty-five days after the infective meal (sufficient time to obtain mature infection in the salivary glands (Van Den Abbeele et al., 1999; Ravel et al., 2003), surviving tsetse flies were individually subjected to a salivation test. Each fly was allowed to salivate into a drop of phosphate-buffered saline– glucose (PSG) on warmed slides (37°C) (Burtt, 1946; Gidudu et al., 1995) and immediately placed in individual cages. The salivate was recovered from the slide in 25 μl of sterile water (Figure 2). To identify flies whose saliva carry trypanosomes, DNA was extracted from each collected saliva and analyzed by TBR1/2 PCR as already described (Ravel *et* al., 2003).

Flies with PCR-positive saliva were then grouped in different cages according to the PCR signal intensity (strong, medium or weak) (Figure 2). Flies with PCR-negative saliva and flies whose salivate could not be recovered because flies refused to salivate, were euthanized and dissected for microscopic observation (x400).

### Monitoring and dissection of the tsetse flies

Midguts of all flies found dead during the process were dissected from day 5 (time necessary to observe parasites colonization of the midgut) to day 19 post-infection (pi). From day 20 pi, the salivary glands were also dissected (assuming that no trypanosomes can be found in the salivary glands before this time). All the dissected midguts and salivary glands were examined for trypanosomes by phase contrast microscopy at 400x magnification.

### Infection of mice

At day 39 and 42 pi, each group of flies with PCR positive saliva was fed twice (3 days apart) on the belly of anesthetized female BALB/c mice previously immuno-suppressed with 0.15mL of cyclophosphamide (ENDOXAN, 20mg/mL) injected subcutaneously (Figure 2). A different mouse was assigned to each group of flies. The objective of this sorting was to maximize the success of infection of one of the mice. Tsetse were starved for three days prior to the mice bloodmeal to increase sting probability. The parasitemia of the mice was then determined daily by microscopy using the rapid “matching” method (Herbert and Lumsden, 1976) on drop of blood collected from the tail of each mouse (Figure 2).

The mice-fed surviving flies were euthanized and dissected for microscopy (x400) observation of midgut and salivary glands.

## Results and Discussion

### Molecular screening of tsetse flies with trypanosomes in their salivary glands

Thirty-five days after tsetse flies’ infection, 80 flies out of 165 were still alive (Figure 3). Out of them, 78 were tested for the detection of trypanosomes in saliva by PCR and 43 (55%) showed a PCR-positive saliva. Out of these 43 flies, 9, 12 and 22 exhibited a strong, medium and weak PCR signal respectively (Supplementary file 1). Between the beginning of the collection of the saliva and the results of the PCR analysis, 8 flies died. The remaining 35 flies were grouped into four cages: two containing each 9 flies with weak PCR signal, one containing 11 flies with medium PCR signal and one containing 6 flies with strong PCR signal (Figure 3).

**Figure 3:**
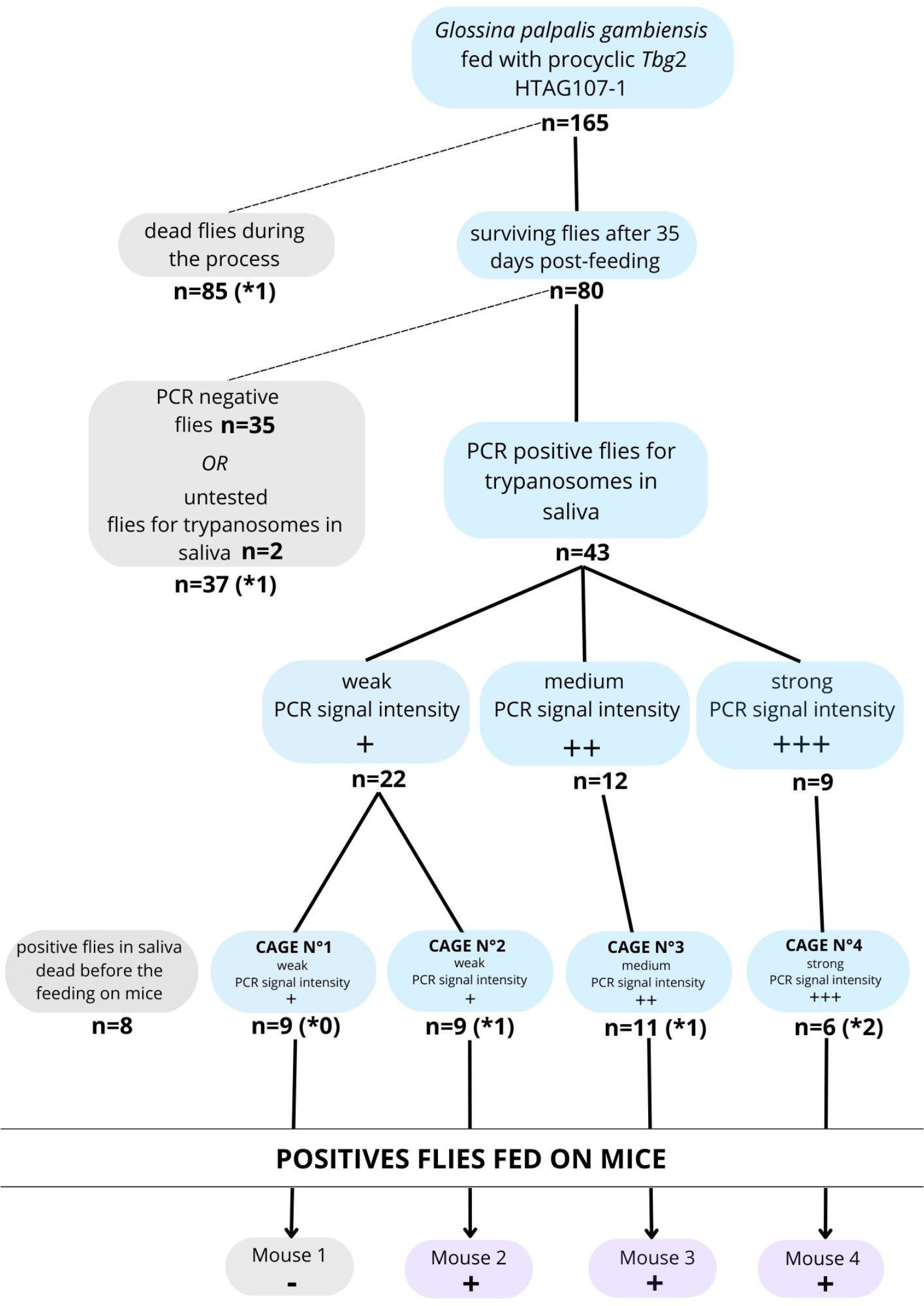
Diagram summarizing the main results of the experimental infection of flies and mice with *Tbg*2 HTAG-107 strain. (*x: number of flies found positive in salivary glands by microscopic observation after dissection).

All the PCR-negative flies were euthanized and dissected for microscopic observation of midgut and salivary glands.

### Monitoring of tsetse flies’ infection

Throughout this experiment, 115 flies could be dissected of which 11 (9.6%) showed parasites in their midgut only and six (5.2%) in both their midgut and salivary glands. Among the flies showing trypanosomes in their salivary glands that were fed on mice, none have been found in cage n°1, one was from cage n°2, one from cage n°3 and two from cage n°4, in line with the molecular analysis.

The number of mature infections identified by the PCR analysis of the flies’ saliva (n=43) was much higher than that determined by microscopic observation of the salivary glands (n=6). This data is in the range with previous observations from other studies (with maximum 10% flies found positives in SG) (Ravel et al., 2003, Ravel et al., 2006; Richner et al., 1988). Because of its high sensitivity, PCR from saliva may offer a better view of mature infections as it is sometimes challenging to observe trypanosomes in salivary glands. However, for most of the PCR positives flies for trypanosomes in their saliva, no trypanosomes were observed in the dissected salivary glands despite diligent microscopic observation.

Additionally, because the recovery of salivary glands is not so easy during the dissection of the flies, salivary glands may not always be available for observation. This was the case for three of the flies dissected at day 36 pi, trypanosomes were observed in the midgut only but the salivary glands could not be recovered for technical reasons. Therefore, the percentage of flies with mature infection may have been underestimated through microscopy techniques.

### Monitoring the infection in mice

Once the trypanosome PCR positive flies of the four cages have been fed on mice, the parasitaemia has been daily monitorized. Three mice out of the four developed an infection two to four days after the blood meal (Figure 3 and supplementary file 2). The resulting HTAG107-1 BSF trypanosomes were collected from the infected mice by cardiac puncture and supplemented with 50% glycerol before being stored in liquid nitrogen. Dissection and observation of the salivary glands of the flies used to bite the mice showed that two flies were positive for trypanosomes in cage n°4, one in cage n°3 and one in cage n°2. No salivary glands positive flies were detected in cage n°1, which is congruent with the absence of infection in mouse 1.

### Advantages and drawbacks to reconstitute a PCF to BSF experimental *in vivo* life cycle

From 165 flies fed with a single meal of sheep blood mixed with cultivated PCF trypanosomes, at least 6 flies with mature infections after one month have been obtained.

We succeeded in infecting mice from infected tsetse flies. This achievement is partly due to the large size of the starting sample (n=165) and to the starvation of the flies before the infective meal. Post-dissection of the flies used to infect the mice showed that a mouse only needs to be stung by one fly with mature infection to become infected. This success also confirms that *Tbg*2 group strains can settle in the tsetse fly and infect a mammalian host.

For strains only available as PCF, result from this experiment makes possible to obtain bloodstream form that can be evaluated for resistance to NHS. However, this experiment is time consuming, requires a great technical effort and is not in line with current animal ethical principles (3R rule (replace, reduce, refine)). For this reason, if the passage from PCF to BSF is the only result desired, the *in vitro* plasmid methods should be preferred (Mugo *et* al. 2017).

While HAT elimination seems a realistic goal for 2030, we advocate for improving the knowledge of the *T. b. gambiense* strains that are still circulating, even at a low level, almost undetectable level. In several HAT foci, parasitaemia observed in human or in animal reservoir is very low (Camara et al., 2010) and hinders a deep genotypical and phenotypical characterization. Isolating strains and mastering a transmission cycle allow to collect data that are not possible to collect otherwise like to account for a difference in pathogenicity and virulence to human over natural cycles or not (Büscher et al., 2018). This is particularly interesting in the case of *Tbg*2 strains. They have been found resistant or partially resistant towards the NHS but their ability to maintain the NHS resistance capacity after cycling in animals is unknown. Deciphering the nature of the resistance to NHS: constitutive (as it is the case for *Tbg*1) or conditional (which can be lost after several vector/animal cycles) is a keystone data for sleeping sickness effective and sustainable elimination. Indeed, elimination of HAT will lead to a decrease in acquired immunity in populations, which could create major concerns for more susceptible populations if they are exposed again to strains with constitutive resistance (Jamonneau et al., 2019).

We provide here the first evidence of an effective *in vivo* protocol to transform *Tbg*2 PCF to BSF by experimentally reproducing the complete *Tbg*2 cycle from *G. p. gambiensis* to the mouse.

## Acknowledgments

Paola JUBAN was granted by KIM RIVE project (Key Initiative Muse Risks & Vectors) from Montpellier Université d’excellence (MUSE).

All experiments on *G. p. gambiensis* have been performed on the Baillarguet insectarium platform (https://doi.org/10.18167/infrastructure/00001), member of the National Infrastructure EMERG’IN and of the Vectopole Sud network (http://www.vectopole-sud.fr/). The Baillarguet insectarium plateform is led by the joint units Intertryp (IRD, Cirad) and ASTRE (Cirad, INRAE).

Mice were kept under strict ethical conditions according to the international guidelines for the care and use of laboratory animals. The experiments designed for this study were approved by the regional Ethic Committee for Animal Experimentation CEEA-LR 36 under project number APAFiS #34149 and authorized by French Ministry for Higher Education and Research.

Figure 1, 2 and Supplementary file 2 are original figures created with Biorender.com.

We would like to thank Bernadette Tchicaya and Annick Ranaivoarisoa for providing males from *G. p. gambiensis*.

