## Supplementary data for "*Trypanosoma brucei gambiense* group 2 experimental *in vivo* life cycle: from procyclic to bloodstream form"

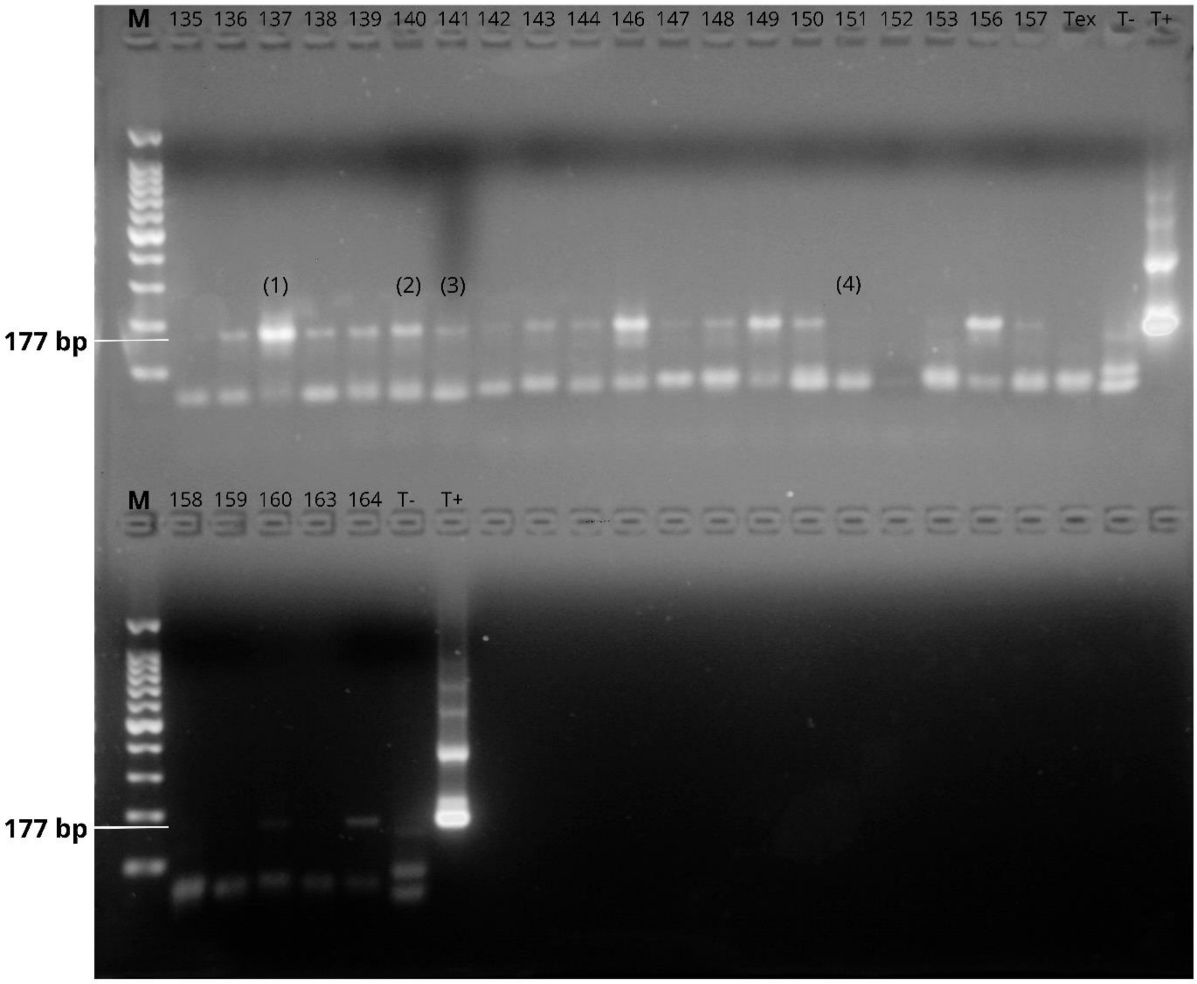


*Supplementary file 1:* TBR1/2 PCR identification of *Trypanosoma brucei s.l* in tsetse flies' saliva showing 177bp DNA satellite repeat specific for *T. brucei s.l*– numbers correspond to those assigned to the flies. M: 100 bp DNA size marker; Tex: DNA extraction negative control; T-: PCR negative control; T+: positive control; (1): Strong PCR signal; (2): Medium PCR signal; (3): Weak PCR signal; (4): Negative PCR signal.

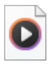

additional figure 2.mp4

*Supplementary file 2:* Microscopic observations videos of *Tbg*2 throughout its life cycle including (1) procyclic form in tsetse midgut, (2) Metacyclic form in tsetse salivary glands and (3) bloostream form found in infected mice blood.
